# Recombinant rotaviruses rescued by reverse genetics reveal the role of NSP5 hyperphosphorylation in the assembly of viral factories

**DOI:** 10.1101/660217

**Authors:** Guido Papa, Luca Venditti, Francesca Arnoldi, Elisabeth M. Schraner, Christiaan Potgieter, Alexander Borodavka, Catherine Eichwald, Oscar R. Burrone

## Abstract

Rotavirus (RV) replicates in round-shaped cytoplasmic viral factories although how they assemble remains unknown.

During RV infection, NSP5 undergoes hyperphosphorylation, which is primed by the phosphorylation of a single serine residue. The role of this post-translational modification in the formation of viroplasms and its impact on the virus replication remains obscure. Here we investigated the role of NSP5 during RV infection by taking advantage of a modified fully tractable reverse genetics system. An NSP5 trans-complementing cell line was used to generate and characterise several recombinant rotaviruses (rRVs) with mutations in NSP5. We demonstrate that a rRV lacking NSP5, was completely unable to assemble viroplasms and to replicate, confirming its pivotal role in rotavirus replication.

A number of mutants with impaired NSP5 phosphorylation were generated to further interrogate the function of this post-translational modification in the assembly of replication-competent viroplasms. We showed that the rRV mutant strains exhibit impaired viral replication and the ability to assemble round-shaped viroplasms in MA104 cells. Furthermore, we have investigated the mechanism of NSP5 hyper-phosphorylation during RV infection using NSP5 phosphorylation-negative rRV strains, as well as MA104-derived stable transfectant cell lines expressing either wt NSP5 or selected NSP5 deletion mutants. Our results indicate that NSP5 hyper-phosphorylation is a crucial step for the assembly of round-shaped viroplasms, highlighting the key role of the C-terminal tail of NSP5 in the formation of replication-competent viral factories. Such a complex NSP5 phosphorylation cascade may serve as a paradigm for the assembly of functional viral factories in other RNA viruses.

**IMPORTANCE:** Rotavirus (RV) double-stranded RNA genome is replicated and packaged into virus progeny in cytoplasmic structures termed viroplasms. The non-structural protein NSP5, which undergoes a complex hyperphosphorylation process during RV infection, is required for the formation of these virus-induced organelles. However, its roles in viroplasm formation and RV replication have never been directly assessed due to the lack of a fully tractable reverse genetics (RG) system for rotaviruses. Here we show a novel application of a recently developed RG system by establishing a stable trans-complementing NSP5-producing cell line required to rescue rotaviruses with mutations in NSP5. This approach allowed us to provide the first direct evidence of the pivotal role of this protein during RV replication. Furthermore, using recombinant RV mutants we shed light on the molecular mechanism of NSP5 hyperphosphorylation during infection and its involvement in the assembly and maturation of replication-competent viroplasms.

## INTRODUCTION

Rotavirus (RV) is the most common cause of viral gastroenteritis in young children and infants worldwide (1, 2). It is a non-enveloped RNA virus with a genome composed of 11 segments of double-stranded RNA (dsRNA), which replicates in cytoplasmic structures primarily composed of viral proteins (3–5). During infection, the first steps of viral morphogenesis and genome replication occur within cytoplasmic viral replication factories known as viroplasms. (3, 5–7). The assembly of viroplasms requires co-expression of at least two non-structural proteins, NSP5 and NSP2 (8, 9), however, how these virus-induced organelles are formed remains unknown.

Other viral proteins also found in viroplasms include RNA-dependent RNA polymerase (RdRp) VP1, the main inner-core protein VP2, guanyltransferase/ methylase VP3, and the middle layer (inner capsid) protein VP6 (10, 11). Biochemical evidence suggests that viroplasms are essential for RV replication since the virus production is highly impaired upon silencing of either NSP2 or NSP5 (12–15).

Rotavirus NSP5, encoded by genome segment 11, is a small serine (Ser)- and threonine (Thr)-rich non-structural protein that undergoes multiple post-translational modifications in virus-infected cells, including O-linked glycosylation (16), N-acetylation (17), SUMOylation (18) and crucially hyperphosphorylation that involves several distinct Ser residues (19, 20). The NSP5 hyperphosphorylation is a complex process, which gives rise to multiple phosphorylation states ranging from the most abundant 28 kDa phospho-isoform, up to the hyperphosphorylated 32-34 kDa states (19, 20). All these forms have been found to be more stable in viroplasms, while chemical disruption of viroplasms results in NSP5 de-phosphorylation (21). The mechanism of NSP5 phosphorylation is not yet wholly understood, but it involves interactions with other viral proteins. When expressed alone in non-infected cells, NSP5 is not phosphorylated, while co-expression with NSP2 or VP2 results in NSP5 hyperphosphorylation and formation of viroplasm-like structures (VLS) (8, 22, 23). NSP5 hyperphosphorylation involves the phosphorylation of Serine 67 (Ser67) by Casein Kinase 1α (CK1α) to initiate the phosphorylation cascade (24, 25) and it is considered to be essential for the assembly of viroplasms (26). Although the structure of NSP5 remains unknown, it readily forms higher molecular weight oligomeric species in solution, potentially providing a larger interface for interacting with multiple components of viroplasms (27).

In addition, the C-terminal region (‘a tail’ including amino acids 180-198) is required for NSP5 decamerisation *in vitro* (27) and VLS formation *in vivo* (7). However, due to the lack of a fully tractable reverse genetics (RG) system for RVs until recently, previous studies on NSP5 have been carried out using the mutants expressed in the absence of a complete set of viral proteins. Here, we took advantage of the novel plasmid only-based, helper-virus free RG systems for rotaviruses (28, 29) to gain new insights into the mechanisms of NSP5 hyperphosphorylation and its role in viroplasm assembly and virus replication during viral infection.

To achieve this, we generated and characterised several viable recombinant rotaviruses (rRVs) with mutations in NSP5. Using these mutants, we show the role of NSP5 hyperphosphorylation for viroplasm assembly and in genome replication. These studies shed light on a complex hierarchical mechanism of NSP5 hyperphosphorylation during rotaviral infection.

## MATERIALS AND METHODS

### Cells and viruses

MA104 (embryonic African green monkey kidney cells, ATCC CRL-2378.1 from *Chlorocebus aethiops*), U2OS (Human bone osteosarcoma epithelial cells), Caco-2 (colorectal adenocarcinoma human intestinal epithelial cell line, ATCC®HTB-37) and HEK293T (embryonic human kidney epithelial, ATCC®CRL-3216) cells were cultured in Dulbecco’s Modified Eagle’s Medium (DMEM) (Life Technologies) supplemented with 10% Fetal Bovine Serum (FBS) (Life Technologies) and 50 μg/ml gentamycin (Biochrom AG).

MA104-NSP5-EGFP cells (MA-NSP5-EGFP) (7) were cultured in DMEM supplemented with 10% FBS (Life Technologies), 50 μg/ml gentamycin (Biochrom AG) and 1 mg/ml geneticin (Gibco-BRL, Life Technologies).

MA104-NSP2-mCherry (MA-NSP2-mCherry), MA104-Δ3 (MA-Δ3), MA104-Δtail (MA-ΔT) and MA104-NSP5wt (MA-NSP5) stable transfectant cell lines (embryonic African green monkey kidney cells, ATCC® CRL-2378) were grown in DMEM (Life Technologies) containing 10% FBS, 50 μg/ml gentamycin (Biochrom AG) and 5 μg/ml puromycin (Sigma-Aldrich).

BHK-T7 cells (Baby hamster kidney stably expressing T7 RNA polymerase) were cultured in Glasgow medium supplemented with 5% FBS, 10% Tryptose Phosphate Broth (TPB) (Sigma-Aldrich), 50 μg/ml gentamycin (Biochrom AG), 2% Non-Essential Amino Acid (NEAA), 1% Glutamine.

Recombinant simian RV strain SA11 (rRV-wt), rescued using reverse genetics system using cDNA clones encoding the wild-type SA11 (G3P[2]) virus (28), was propagated in MA104 cells cultured in DMEM supplemented with 0.5 μg/ml trypsin (Sigma Aldrich).

### Recombinant RVs Titration

Recombinant NSP5 mutant RVs were grown in MA-NSP5 cells and the lysate was 2-fold serially diluted and used to infect MA-NSP5 cells, seeded in 24-wells plates with coverslips. After 1 hour of adsorption, virus was removed, and cells were incubated at 37°C. At 5 hours post-infection (hpi), cells were fixed with 4% paraformaldehyde (PFA) in phosphate buffer saline (PBS) [137 mM NaCl; 2.7 mM KCl; 8.1 mM Na_2_HPO_4_ and 1.74 mM KH_2_PO_4_ pH7.5] for 15 min at room temperature and permeabilized for 5 min with PBS containing 0.01% Triton X-100. Next, cells were incubated for 30 min with PBS supplemented with 1% bovine serum albumin (PBS-BSA) at room temperature and then with anti-NSP5 (1:1000) or anti-VP2 (1:200) or anti-NSP2 (1:200) guinea pig serum diluted in PBS-BSA. After washing three times with PBS, cells were incubated for 1 h at room temperature with TRITC-conjugated Anti-guinea Pig IgG (Jackson ImmunoResearch) (1:500) diluted in PBS-BSA.

Nuclei were stained with ProLong™ Diamond Antifade Mountant with DAPI (Thermo Scientific). Samples were imaged using a confocal setup (Zeiss Airyscan equipped with a 63x, NA=1.3 objective). Each viroplasms-containing cell was counted as one focus-forming unit (FFU). The average of cells with viroplasms of six fields of view per each virus dilution was determined and the total number of cells containing viroplasms in the whole preparation was estimated. The virus titre was determined as:

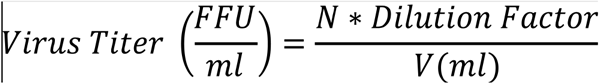

where *N* is a total number of cells containing 1 or more viroplasms, and *V* is the volume of virus inoculum added.

### Replication Kinetics of recombinant viruses

MA104 cell line (ATCC CRL-2378.1) or stably transfected MA104 cells (NSP5; Δ3, ΔT) were seeded into 24-well plates and subsequently infected with recombinant RVs at MOI (FFU/cell) of 0.5 for multi-step growth curve experiments and MOI 5 for a single-step growth curve experiment. After adsorption for 1 h at 37°C, cells were washed twice with PBS and the medium replaced with DMEM without trypsin. After incubation at 37°C, cells were harvested after 8, 16, 24, 36 hours post virus adsorption. The cell lysates were freeze-thawed three times and activated with trypsin (1 μg/ml) for 30 min at 37°C. The lysates were used to infect monolayers of MA-NSP5 cells seeded in µ-Slide 8 Well Chamber Slide-well (iBidi GmbH, Munich, Germany). The cells were then fixed 5 hours post infection for 15 min with 4% paraformaldehyde and permeabilized for 5 min with PBS containing 0.01% Triton X-100. Next, cells were incubated for 30 min with PBS-BSA at room temperature and then with anti-NSP5 serum (1:1000) diluted in PBS-BSA. After three washes with PBS, cells were incubated for 1 h at room temperature with TRITC-conjugated Anti-guinea Pig IgG (Jackson ImmunoResearch) (1:500) diluted in PBS containing 1% BSA (PBS-BSA).

The number of infected cells was counted, and the virus titres were expressed in Focus-Forming Units per mL (FFU/mL).

### Plasmid construction

RV plasmids pT_7_-VP1-SA11, pT_7_-VP2-SA11, pT_7_-VP3-SA11, pT_7_-VP4-SA11, pT_7_-VP6-SA11, pT_7_-VP7-SA11, pT_7_-NSP1-SA11, pT_7_-NSP2-SA11, pT_7_-NSP3-SA11, pT_7_-NSP4-SA11, and pT_7_-NSP5-SA11 (28) were used to rescue recombinant RVs by reverse genetics. pT_7_-NSP5/S67A carrying a mutation in the nucleotide T220G in the gs11 and pT_7_-NSP5/Tyr18Stop harbouring a nucleotide substitution T75G were generated by QuikChange II Site-Directed Mutagenesis (Agilent Technologies). pT_7_-NSP5/ΔT was generated from pT_7_-NSP5-SA11 by deleting the last 18 C-terminal amino acids (FALRMRMKQVAMQLIEDL) using substitution of the F181 encoding triplet with a stop codon. pT_7_-NSP5/Δ176-180 were obtained deleting the amino acids 176 to 180 (YKKKY). The described deletions were performed using the QuikChange II Site-Directed Mutagenesis kit (Agilent Technologies).

For the generation of lentiviral plasmids, NSP5 and NSP2-mCherry were amplified by PCR and inserted into the plasmid pAIP (Addgene #74171; (30) at the *NotI*-*EcoRI* restriction enzymes sites to yield pAIP-NSP5 and pAIP-NSP2-mCherry, which were then used to generate lentiviruses for the MA104-stable transfectant cell lines (MA-NSP5 and MA-NSP2-mCherry) (31). NSP5/ΔT was amplified from the pT_7_-NSP5/ΔT by PCR and inserted into the pPB-MCS (Vector Builder) at restriction enzyme sites *Nhe*I-*Bam*HI to generate pPB-NSP5/ΔT for the production of the stable transfectant MA-ΔT.

pPB-NSP5/Δ3, for MA-Δ3 stable cell line establishment, was generated with a GenParts™ DNA Fragment (GenScript) containing NSP5 ORF lacking the amino acids 80-130 (VKTNADAGVSMDSSAQSRPSSNVGCDQVDFSLNKG LKVKANLDSSISIST) and inserted into the *Nhe*I-*BamH*I restriction sites of pPB-MCS vector.

### Generation of stable cell lines

MA-NSP5-EGFP were generated as previously described (7).

MA-NSP2-mCherry and MA-NSP5 cell line were generated using lentiviral vector system (31). Briefly, HEK293T cells were maintained in DMEM (Life Technologies) supplemented with 10% FBS (Life Technologies), and 50 μg/ml gentamycin (Biochrom AG). Approximately 7 × 10^6^ HEK293T cells were seeded in 10 cm^2^ tissue culture dishes 24 hours before transfection. For each well, 2.4 μg of pMD2-VSV-G, 4 μg of pMDLg pRRE, 1.8 μg of pRSV-Rev and 1.5 μg of plasmid containing pAIP-NSP2-mCherry or pAIP-NSP5 and the human immunodeficiency virus long terminal repeats were co-transfected with Lipofectamine 3000 (Sigma-Aldrich) according to the manufacturer’s instructions. After 48 h, the virus was collected by filtration with a 0.45-μm polyvinylidene fluoride filter and was immediately used or stored at −80 °C. For lentiviral transduction, MA104 cells were transduced in six-well plates with 1 ml of lentiviral supernatant for 2 days.

MA-Δ3 and MA-ΔT were generated using the PiggyBac Technology (32). Briefly, 10^5^ MA104 cells were transfected with the pCMV-HyPBase (32) and the transposon plasmid pPB-NSP5/Δ3 and pPB-NSP5/ΔT using a ratio of 1:2.5 with Lipofectamine 3000 (Sigma-Aldrich) according to the manufacturer’s instructions. The cells were maintained in DMEM supplemented with 10% FBS for 3 days and then the cells were incubated with DMEM supplemented with 10% FBS and 5 μg/ml puromycin (Sigma-Aldrich) for 4 days to allow the selection of cells expressing the gene of interest.

### Rescue of recombinant RVs (rRVs) from cloned cDNAs

To rescue recombinant RV strain SA11 (rRV-WT), monolayers of BHK-T7 cells (4 × 10^5^) cultured in 12-well plates were co-transfected using 2.5 μL of TransIT-LT1 transfection reagent (Mirus) per microgram of DNA plasmid. Each mixture comprised 0.8 μg of SA11 rescue plasmids: pT_7_-VP1, pT_7_-VP2, pT_7_-VP3, pT_7_-VP4, pT_7_-VP6, pT_7_-VP7, pT_7_-NSP1, pT_7_-NSP3, pT_7_-NSP4, and 2.4 μg of pT_7_-NSP2 and pT_7_-NSP5 (29). Furthermore 0.8 μg of pcDNA3-NSP2 and 0.8 μg of pcDNA3-NSP5, encoding NSP2 and NSP5 proteins, were also co-transfected to increase rescue efficiency.

To rescue recombinant rRVs encoding NSP5 mutants, pT_7_ plasmids encoding NSP5/S67A, NSP5/Y18Stop, NSP5/Δ180-198 segments were used instead of pT_7_-NSP5. At 24 h post-transfection, MA-NSP5 cells (5 × 10^4^ cells) were added to transfected cells to provide a functional NSP5 for the virus rescue. The cells were co-cultured for 3 days in FBS-free medium supplemented with trypsin (0.5 μg/mL) (Sigma Aldrich). After incubation, transfected cells were lysed by freeze-thawing and 200 μl of the lysate was transferred to fresh MA-NSP5 cells. After adsorption at 37°C for 1 hour, cells were washed three times with PBS and further cultured at 37°C for 4 days in FBS-free DMEM supplemented with 0.5 μg/mL trypsin (Sigma Aldrich, 9002-07-7) until a clear cytopathic effect was visible. The recombinant viruses were then checked by RT-PCR.

### Immunofluorescence microscopy

Immunofluorescence experiments were performed using µ-Slide 8 Well Chamber Slide-well (iBidi GmbH, Munich, Germany) and the following antibody dilutions: anti-NSP5 guinea pig serum 1:1,000; anti-NSP2 guinea pig serum 1:200; anti-VP2 guinea pig serum 1:500; anti-VP6 mouse monoclonal antibody 1:1,000; Alexa Fluor 488-conjugated anti-mouse, 1:500 (Life Technologies), and TRITC-conjugated anti-guinea pig, 1:500 (Life Technologies).

### 5-Ethynyl-Uridine (EU) labeling

Newly synthesized RNAs were labeled by including 2 mM 5-ethynyl uridine (EU) into the cell culture medium, and modified incorporated nucleotides were reacted with an azide-conjugated fluorophore Alexa-488 following the manufacturer’s protocol for Click-iT RNA Alexa Fluor 488 imaging kit (Thermo Fisher Scientific). Cell nuclei were stained with ProLong™ Diamond Antifade Mountant with 4’,6-diamidino-2-phenylindole (DAPI, Thermo Scientific). Samples were imaged using a confocal setup (Zeiss Airyscan equipped with a 63x, NA=1.3 objective), and the images were processed using ZEN lite software.

### RNA Fluorescence in situ Hybridization (FISH)

Rotavirus-infected MA104 cells were fixed with 4% (v/v) paraformaldehyde in nuclease-free Dulbecco’s phosphate saline buffer (DPBS) for 10 min at room temperature. Samples were washed twice with DPBS, and then permeabilized with 70% (v/v) ethanol in RNAse-free water at +4°C for at least 1 hour prior to hybridization. Permeabilized samples were re-hydrated for 5 min in a pre-hybridization buffer (300 mM NaCl, 30 mM trisodium citrate, pH 7.0 in nuclease-free water, 10 % v/v formamide, supplemented with 2 mM vanadyl ribonucleoside complex). Re-hydrated samples were hybridized with 62.5 nM of an equimolar mixture of Cy3-labelled DNA probes designed to target the coding region of the gene segment 6 of simian rotavirus A/SA11 (Genbank Acc. AY187029.1) using Stellaris Probe Designer v2 software (LCG Biosearch Technologies), in a total volume of 200 µl of the hybridization buffer (Stellaris RNA FISH hybridization buffer, SMF-HB1-10, Biosearch Technologies, supplemented with 10% v/v of deionized formamide). After 4-8 hours of incubation at 37°C in a humidified chamber, samples were briefly rinsed with the wash buffer (300 mM NaCl, 30 mM trisodium citrate, pH 7.0, 10 % v/v formamide in nuclease-free water), after which a fresh aliquot of 300 µl of the wash buffer was applied to each sample and incubated twice at 37°C for 30 min. After 2 washes, nuclei were briefly stained with 300 nM DAPI solution in 300 mM NaCl, 30 mM trisodium citrate, pH 7.0, and the samples were finally rinsed with and stored in the same buffer without DAPI prior to imaging.

### Transmission electron microscopy

MA104 cells were seeded at 1×10^5^ cells in 2 cm^2^ wells onto sapphire discs and infected at MOI of 75 FFU/cell. At 10 hpi, cells were fixed with 2.5% glutaraldehyde in 100 mM Na/K phosphate buffer, pH 7.4 for 1 h at 4°C and kept in that buffer overnight at 4°C. Afterward, samples were postfixed with 1% osmium tetroxide in 100 mM Na/K phosphate buffer for 1h at 4°C and dehydrated in a graded ethanol series starting at 70%, followed by two changes in acetone, and embedded in Epon. Ultrathin sections (60 to 80 nm) were cut and stained with uranyl acetate and lead citrate (33). Samples were analysed in a transmission electron microscope (CM12; Philips, Eindhoven, The Netherlands) equipped with a charge-coupled-device (CCD) camera (Ultrascan 1000; Gatan, Pleasanton, CA, USA) at an acceleration of 100 kV.

### λ-Protein phosphatase and Calf Intestinal Alkaline Phosphatase Assays

Cellular extracts were incubated with 2,000 units of λ-protein phosphatase (λ-Ppase) (New England Biolabs) in 50 mM Tris-HCl (pH 7.5), 0.1 mM EDTA, 5 mM DTT, 0.01% Brij 35, and 2 mM MnCl_2_. The mixture was incubated at 30°C for 2 h. Samples were loaded in SDS-PAGE and analysed by Western blotting.

For the Calf Intestinal Alkaline Ppase (CIP) assay, cellular extracts were incubated with 1,000 units of CIP (New England Biolabs) in CutSmart™ reaction buffer (New England Biolabs) and incubated at 30°C for 2h. Samples were subjected to SDS-PAGE and analysed by Western blotting.

### Electrophoresis of Viral dsRNA Genomes

Cells were infected with the recombinant viruses at MOI of 5 and were lysed 16 hours post infection. Total RNA was extracted from lysed cells with RnaZol*^®^* (Sigma-Aldrich) according the manufacturer’s protocol and the dsRNA segments were resolved on 10% (wt/vol) poly-acrylamide gels (PAGE) for 2 hours at 180 Volts and visualized by ethidium bromide staining (1μg/ml).

### Protein Analysis

Proteins derived from rRVs-infected cellular extract were separated on an SDS-PAGE for 2 hours at 35 mA and transferred to polyvinylidene difluoride membranes (Millipore; IPVH00010) for 1 hour and 30 minutes at 300 mA (24). For protein analysis membranes were incubated with the following primary antibodies: anti-NSP5 (1:5,000) (16), anti-VP2 (1:5,000) (10), anti-NSP2 guinea pig sera (1:2,000). The membranes were then incubated with the corresponding horseradish peroxidase (HRP)-conjugated goat anti-guinea pig (1:10,000; Jackson ImmunoResearch). Mouse HRP-conjugated anti-actin mAb (1:35,000) (clone AC-15, Sigma-Aldrich) was used as loading control.

Signals were detected using the enhanced chemiluminescence system (Pierce ECL Western blotting substrate; Thermo Scientific).

### Statistics used

Statistical analysis and plotting were performed using GraphPad Prism 6 software (GraphPad Prism 6.0, GraphPad Software Inc., La Jolla, CA, USA). Error bars represent standard deviation. Data were considered to be statistically significant when *p* < 0.05 by Student’s t test.

## RESULTS

### Generation of gs11 mutated recombinant RVs

Recombinant rotaviruses (simian rotavirus A strain SA11) carrying different mutations in the NSP5 coding region of gs11 were obtained using a recently developed reverse genetics protocol (29). Two essential additional modifications were introduced to *trans*-complement the potential loss of NSP5 function in the rRV mutants: i) an additional T7-driven plasmid encoding the ORF of wt NSP5 (i.e., without gs11 5’ and 3’ UTRs) was included in the transfection step of BHK-T7 cells; and ii) each rescued rRV was amplified in a stable transfectant cell line MA-NSP5, supplying the wt NSP5 in trans. Crucially, we have successfully established a novel MA-NSP5 cell line to support the replication of NSP5-deficient recombinant viruses to enable their further in-depth characterisation. Both steps (i) and (ii) were absolutely required for rescuing the NSP5 KO virus, confirming the essential role of NSP5 for RV replication. These mutants, as well as additional stable cell lines generated for this study as described in Materials and Methods, are summarised in Table 1.

**Table 1.**
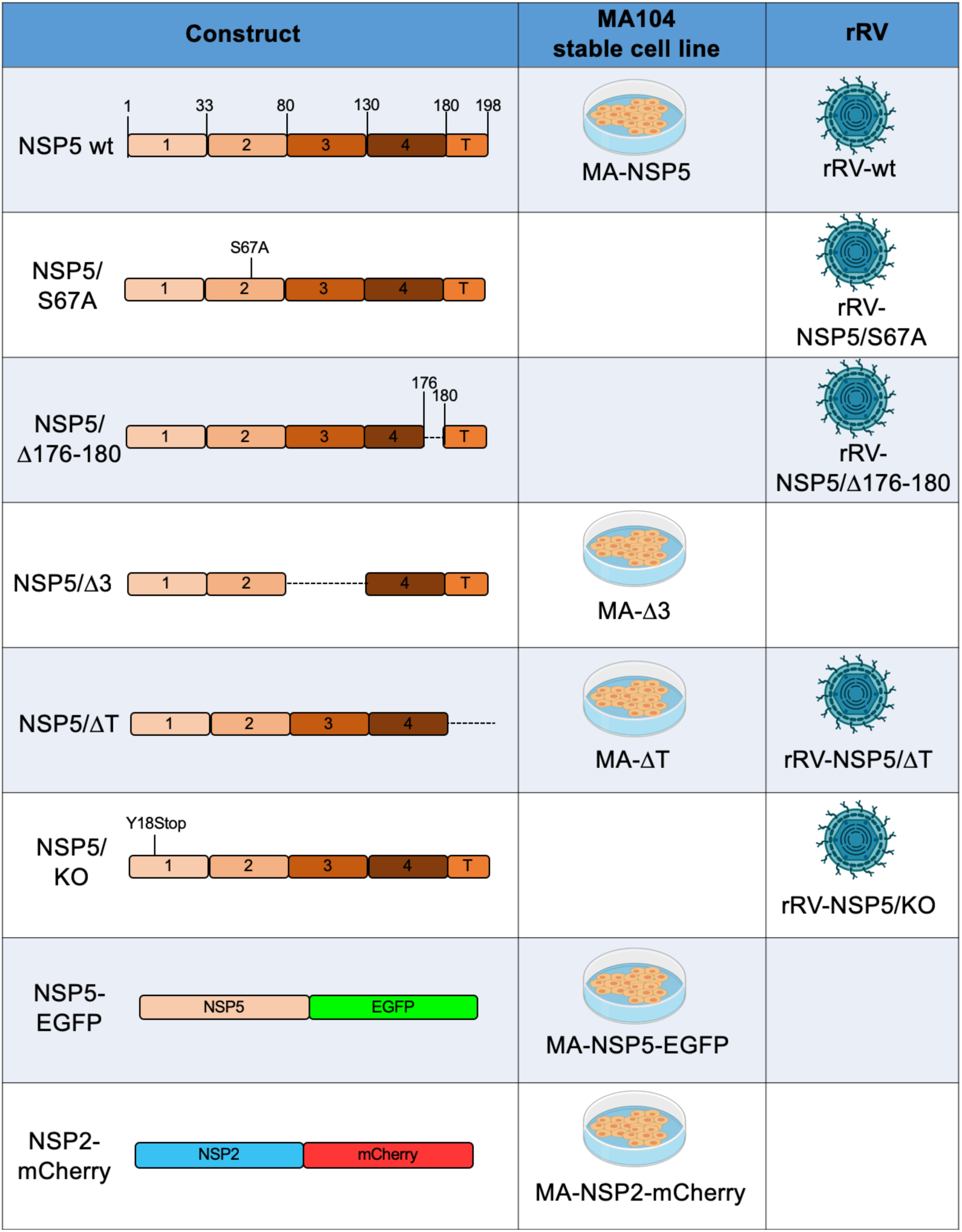
Schematic representation of NSP5 and NSP2 mutants or fusion proteins used to generate rRV and/or stable MA104 transfectant cell lines, used in this study.

| Construct | MA104 stable cell line | rRV |
| --- | --- | --- |
| NSP5 wt | MA-NSP5 | rRV-wt |
| NSP5/<br>S67A |  | rRV-NSP5/S67A |
| NSP5/<br>Δ176-180 |  | rRV-NSP5/Δ176-180 |
| NSP5/Δ3 | MA-Δ3 |  |
| NSP5/ΔT | MA-ΔT | rRV-NSP5/ΔT |
| NSP5/<br>KO |  | rRV-NSP5/KO |
| NSP5-EGFP | MA-NSP5-EGFP |  |
| NSP2-mCherry | MA-NSP2-mCherry |  |

#### rRV-NSP5/KO

NSP5 expression and localisation to viroplasms in virus-infected cells have been considered essential for virus replication (12, 13, 25, 34). Previous studies, using siRNA targeting gs11 mRNA have shown strong impairment of RV replication (12, 13). In order to investigate the effects of point mutations and deletions within the NSP5 gene, we took advantage of the established trans-complementing MA104 cell line stably expressing NSP5. In addition, we also made two cell lines expressing NSP2-mCherry and NSP5-EGFP fusions, which are rapidly and efficiently recruited into viroplasms upon virus infection (Table 1) (7, 22).

Here we provide direct demonstration of the role of NSP5 in RV replication, using an NSP5 knock out rRV (termed rRV-NSP5/KO) generated by reverse genetics. To rescue the NSP5/KO strain, a stop codon at position 18Y was introduced (Fig. 1A-B). Analysis of MA104-virus-infected cell extracts confirmed the presence of NSP2 and VP2, but not of NSP5 (Fig. 2A). Moreover, we did not detect viroplasms containing NSP2, VP2 or VP6 in both MA104-infected cells and in stable transfectant cell lines expressing the fluorescent fusion proteins NSP2-mCherry or NSP5-EGFP (Fig. 2B). Interestingly, this indicates that the NSP5-EGPF fusion protein is not able to trans-complement the lack of NSP5, and indeed the rRV-NSP5/KO strain does not replicate in MA-NSP5-EGFP cells. Furthermore, both genomic dsRNA synthesis and infectious progeny virus production were completely abrogated in MA104 cells, but not in the trans-complementing MA-NSP5 cell line (Fig. 2C). Together, these data confirm that NSP5 is essential for RV replication.

**Figure 1.**
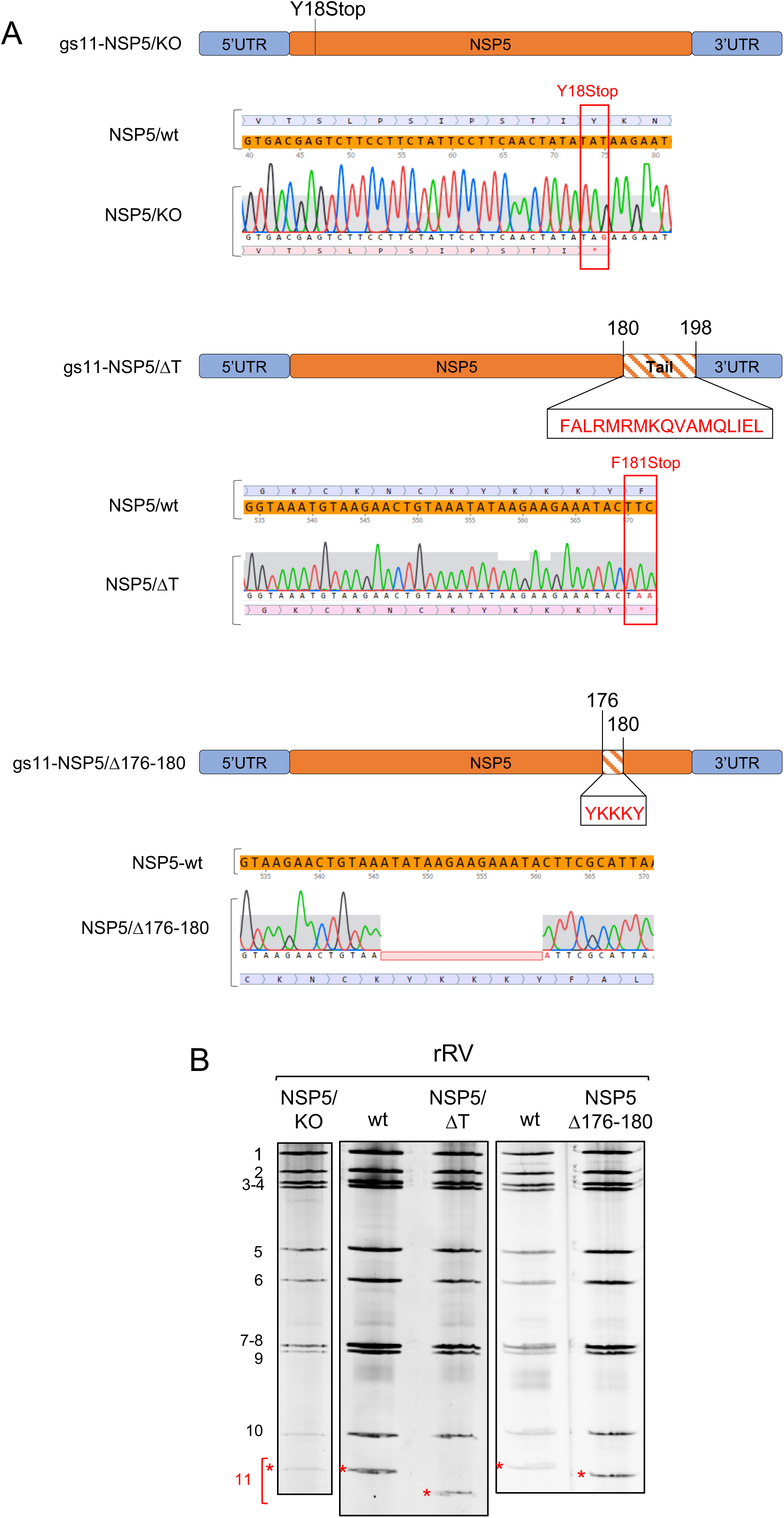
A) Schematic representation of mutations in gs11 of the corresponding rRV strains. Sequences mutated or deleted in NSP5 are indicated. B), Profiles of viral dsRNAs of the different rRV strains generated grown in MA-NSP5 cells. gs11, indicated in red with *.

**Figure 2.**
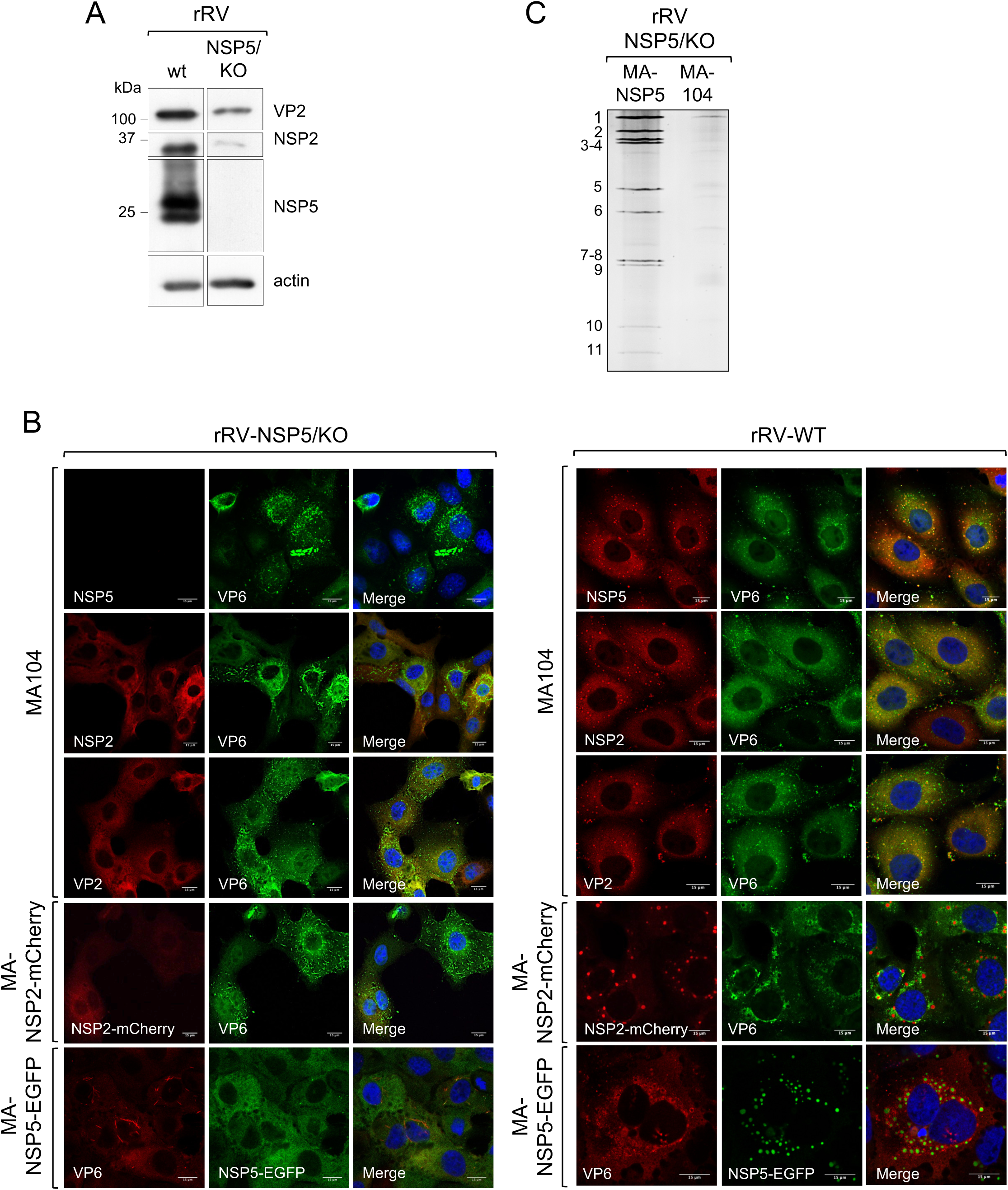
Characterisation of rRV-NSP5/KO. A) Western blot of MA104 cells extracts infected with rRV-wt and rRV-NSP5/KO strains (MOI of 1 FFU/cell). B) Confocal immunofluorescence microscopy of MA104, MA-NSP2-mCherry and MA-NSP5-EGFP cells infected with rRV-wt or rRV-NSP5/KO (MOI of 1 FFU/cell), using antibodies for NSP5, NSP2, VP2 and VP6, as indicated. Scale bar, 15 μm. C) Electrophoretic migration pattern of dsRNAs extracted from rRV-NSP5/KO strain grown in MA-NSP5 or MA104 cells. Genome segments 1-11 are indicated to the left.

#### rRVs with impaired NSP5 phosphorylation

To address the role of NSP5 hyperphosphorylation, we then generated a number of rRVs harbouring NSP5 mutations previously known to impact NSP5 phosphorylation. We first generated an rRV carrying an S67A mutation (rRV-NSP5/S67A) (Table 1 and Fig. 3A-B). Its replication in wild type MA104 cells was strongly impaired (Fig. 3B right panel), resulting in approximately a 100-fold reduction of the infectious progeny virus titre at different time points post-infection (Fig. 3C). Despite the overall reduction of replication fitness, the rRV-NSP5/S67A mutant virus was stable after 10 passages in the wild-type MA104 cells, confirmed by the sequencing of the progeny virus. Consistent with our previous results, the NSP5-S67A mutant was not hyper-phosphorylated in all cell lines tested, including MA104, U2OS and Caco-2 cells (Fig. 3D), further confirming the role of Ser67 in the initiation of NSP5 phosphorylation cascade (24). While the wt rRV yielded multiple hyper-phosphorylated NSP5 isoforms, the NSP5/S67A mutant mostly produced a single, homogeneous form of NSP5 with an apparent mass of 26 kDa that could be detected in the virus-infected cell extracts at 5 or 10 hpi (Fig. 3E). Enzymatic de-phosphorylation with λ-Ppase and alkaline Calf Intestinal Ppase (CIP), previously used to discriminate phosphorylated from non-phosphorylated NSP5 (19), further corroborated the observed lack of NSP5/S67A phosphorylation (Fig. 3F). Because of the differences in the molecular weight markers used, the NSP5 band with the fastest PAGE mobility has been traditionally described as 26 kDa, and the most abundant one as 28 kDa. Since this nomenclature has been used in many publications, we have preferred to maintain it, despite current PAGE migrations do not correspond to the markers presently used.

**Figure 3.**
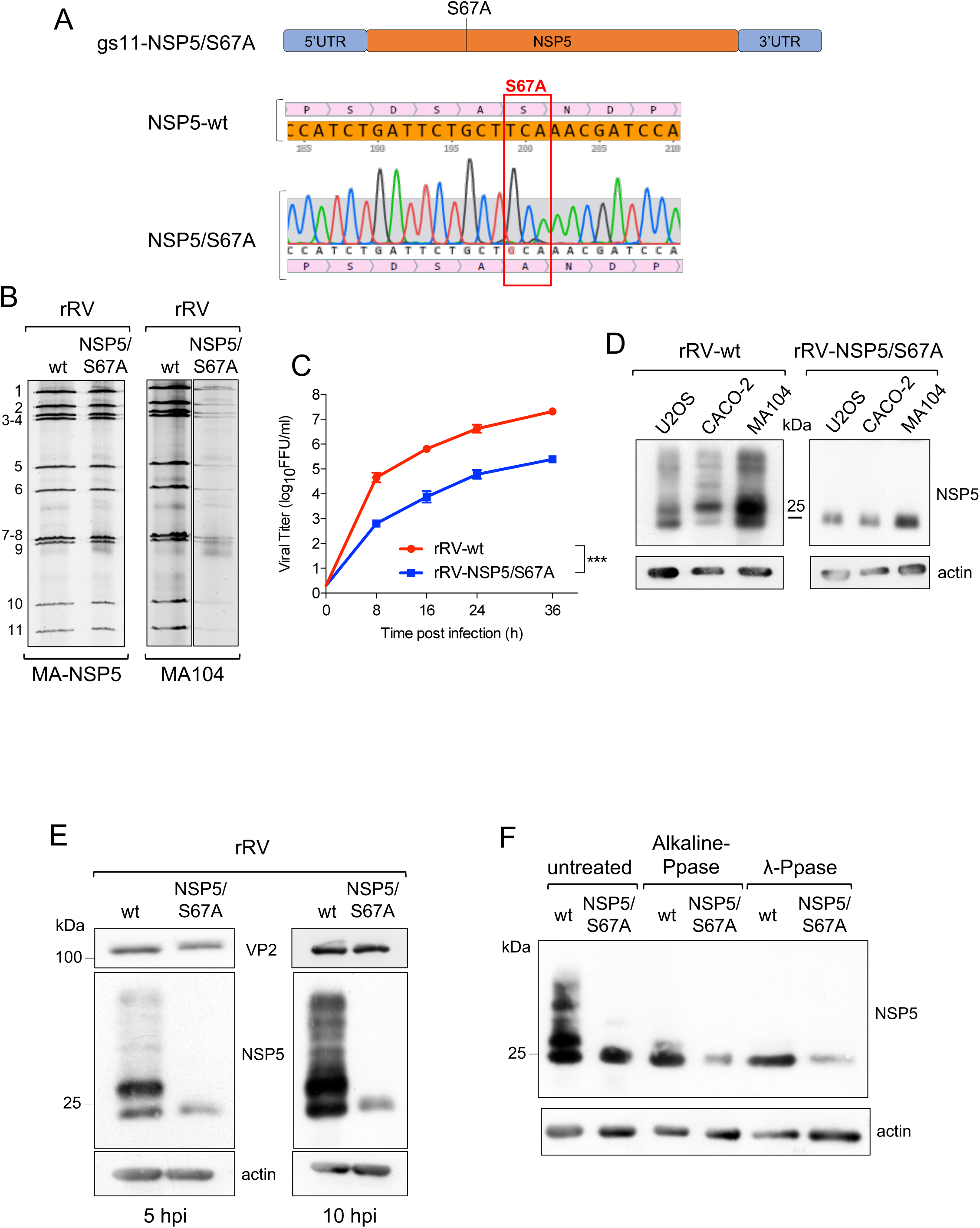
Characterisation of rRV-NSP5/S67A. A) Schematic representation of rRV-NSP5/S67A gs11 and sequence of NSP5 wt and S67A mutant (highlighted). B) Electrophoretic pattern of dsRNA genome segments of rRV-wt and rRV-NSP5/S67A strains grown in MA-NSP5 cells (left panel) and in MA104 wt cells (right panel). C) Replication kinetics of rRV-wt and rRV-NSP5/S67A in MA104 cells. Data are expressed as the means +/- standard deviations (n=3); ***, *p* < 0.001 (Student’s t test). D) Western blot of extracts of U2OS, Caco-2 and MA104 cells infected with rRV-wt or rRV-NSP5/S67A strains. E) Western blot of NSP5 phosphorylation pattern in MA104 cells infected with rRV-wt or rRV-NSP5/S67A (MOI of 1 FFU/cell) at 5 and 10 hpi. F) Western blot of λ-Ppase and alkaline Ppase treatment of lysates of MA104 cells infected with rRV-wt or rRV-NSP5/S67A. Protein bands corresponding to NSP5 are shown.

We then generated two additional rRVs harbouring truncated versions of NSP5, one lacking the 18 AA long C-terminal tail (rRV-NSP5/ΔT), and the second one with a 5 AA deletion (176-YKKKY-180) just upstream of the tail region (rRV-NSP5/Δ176-180) (Table 1 and Fig. 1A-B). Despite the presence of Ser67, and the lack of Thr and Ser residues within the deletions, both mutants did not show a classical hyperphosphorylation pattern (Fig. 4A) and failed to replicate, confirmed by the absence of *de novo* synthesis of genomic dsRNA (Fig. 4B).

**Figure 4.**
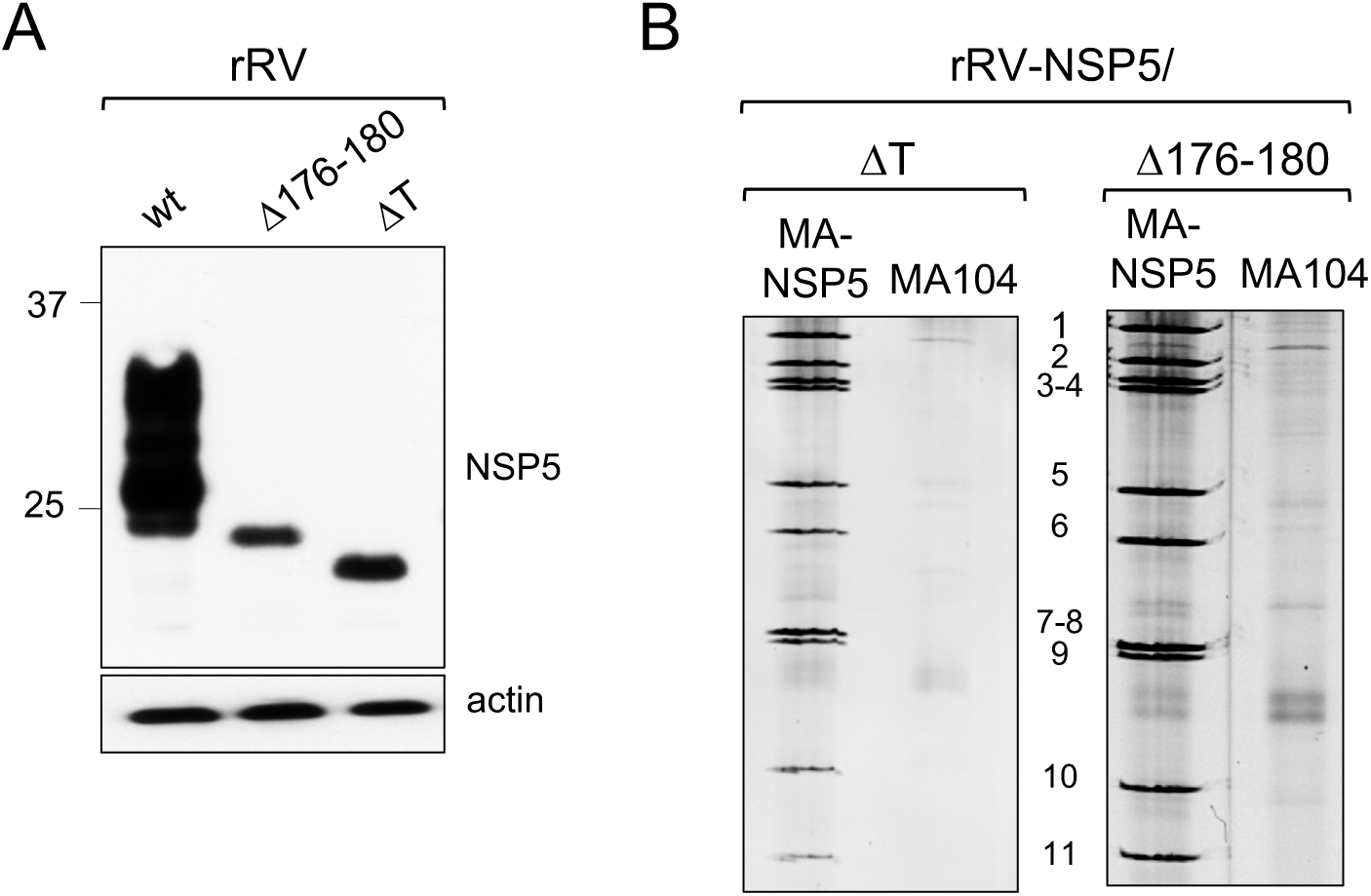
Characterisation of rRV-NSP5/ΔT and rRV-NSP5/Δ176-180. A) Western blot of NSP5 and NSP5 mutants from MA104 cells infected with rRV-wt, rRV-NSP5/ΔT or rRV-NSP5/Δ176-180 at 5 hpi. B) Electrophoretic dsRNA migration pattern of rRV-NSP5/ΔT and rRV-NSP5/Δ176-180 grown in MA-NSP5 and MA104 cells.

We also investigated viroplasm formation in cells infected with these rRV mutants. At early infection (5 hpi), the rRV-NSP5/S67A mutant produced structures resembling viroplasms that appeared smaller and more heterogeneous in shape compared to the regular, spherical ones produced during the wt rRV infection (Fig. 5A upper panel and Fig. 5B). Remarkably, during late infection (10-12 hpi), the rRV-NSP5/S67A mutant produced multiple NSP5-containing aberrant structures, as well as fibre-like structures that became more apparent during late infection (12 hpi) (Fig. 5A lower panel and Fig. 5B).

**Figure 5.**
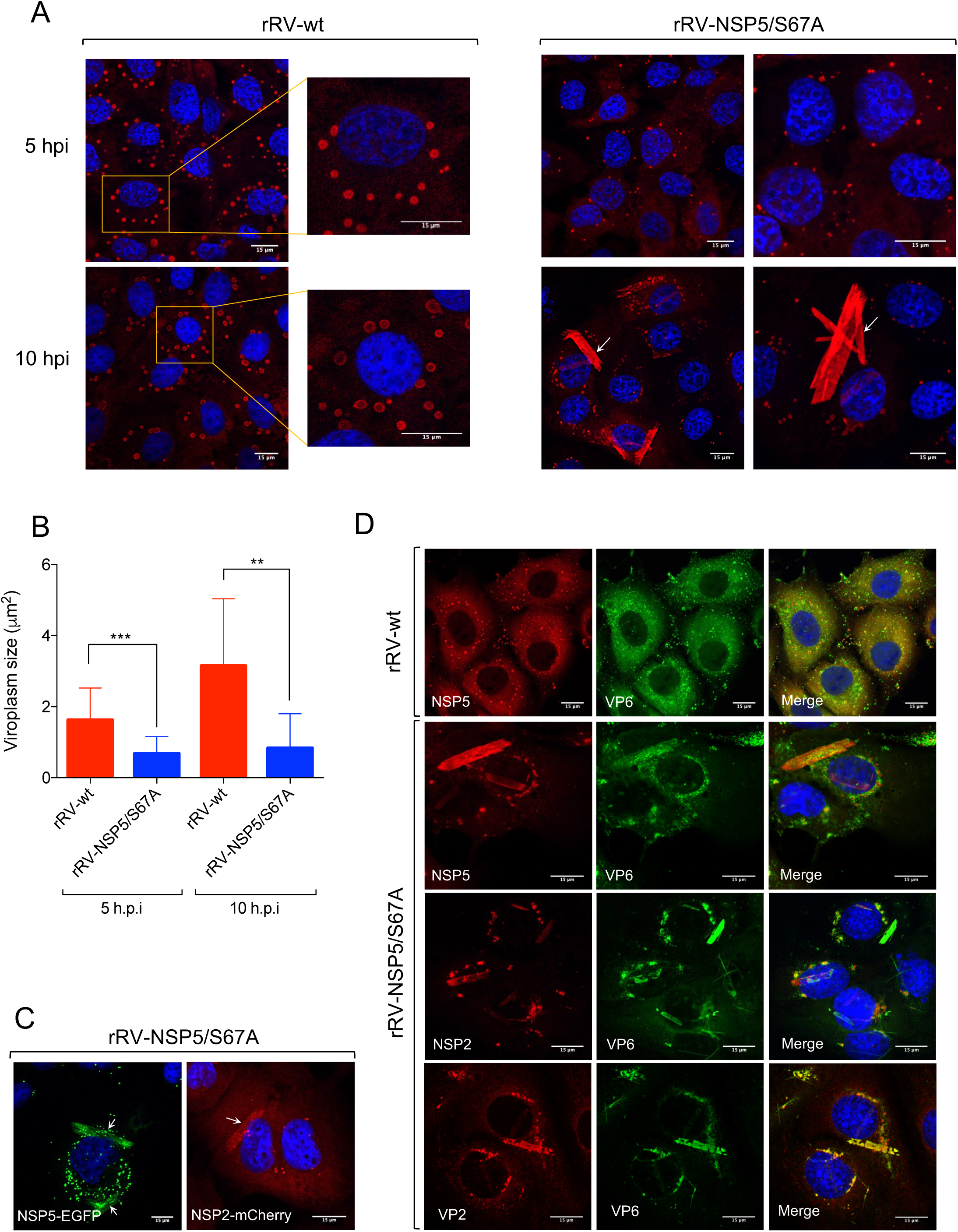
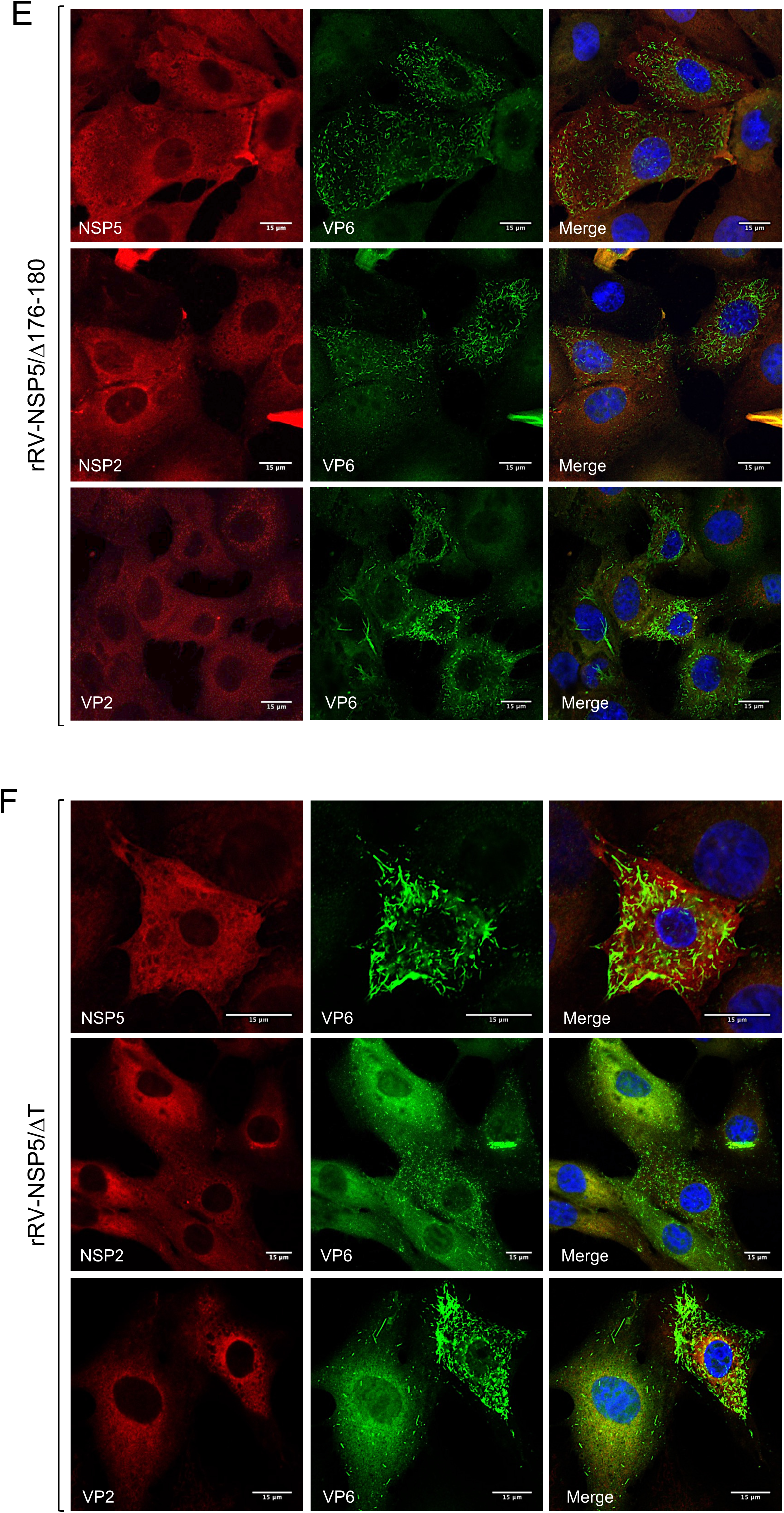
Viroplasms morphology in MA104 cells infected with rRV-NSP5 wt and mutants. Representative confocal immunofluorescence micrographs of MA104 cells infected with rRV-wt and rRV-NSP5/S67A (MOI of 15 FFU/cell) at 5 hpi (upper panels) or 10 hpi (lower panels). Cells were stained with anti-NSP5 and DAPI. B) Quantitative analysis of viroplasms size (μm^2^) of MA104 cells infected with rRV-wt and rRV-NSP5/S67A at 5 hpi and 10 hpi. ***, p < 0.001; **, p < 0.01. C) Confocal immunofluorescence micrographs of MA-NSP5-EGFP and MA-NSP2-mCherry cells infected with rRV-NSP5/S67A (MOI of 15 FFU/cell). D) Confocal immunofluorescence micrographs of rRV-wt and rRV-NSP5/S67A infected MA104 cells (MOI 15 FFU/cell) and stained at 10 hpi with the indicated antibodies. E), F), Confocal immunofluorescence of MA104 cells infected with rRV-NSP5/Δ176-180 (E) or rRV-NSP5/ΔT (F) and stained with the indicated antibodies and DAPI. Scale bar, 15 μm.

Interestingly, these structures produced by rRV-NSP5/S67A were morphologically similar to those observed during wt RV infection of MA104 cells silenced for cellular kinase CK1α, previously shown to be required for NSP5 phosphorylation (25). NSP2-mCherry and NSP5-EGFP fusion proteins were also recruited to both types of these structures (Fig. 5C), suggesting that the observed lack of phosphorylation does not affect NSP2/NSP5 interactions. Furthermore, the NSP5/S67A mutant and additional viroplasmic proteins NSP2, VP6 and VP2 (Fig. 5D) were all present in the aberrant structures, suggesting that they could also represent sites of virus replication.

In contrast, no viroplasms containing NSP5, NSP2 or VP2 were observed when MA104 cells were infected with the rRV-NSP5/ΔT and rRV-NSP5/Δ176-180 deletion mutants. Interestingly, these mutants yielded fibre-like structures containing only VP6 protein (Fig. 5E-F). Similar VP6-fibres are normally formed when VP6 is over-expressed in cells in the absence of other viral proteins (35, 36).

Taken together, these results confirm the role of NSP5 hyper-phosphorylation for controlling the assembly of regular-shaped viroplasms, highlighting the key role of the C-terminal tail in the formation of RV viral factories.

#### RNA accumulation in aberrant structures

Having examined the viral protein composition of the aberrant viroplasms formed during infection with rRVs exhibiting impaired NSP5 phosphorylation, we then assessed their RNA content. Viral RNA transcripts were labelled by incorporation of 5-ethynyl uridine (5-EU) in actinomycin D-treated RV-infected cells, and total viral ssRNA was visualized by reacting with Alexa-488-azide, as described in Materials and Methods. As expected, most viral transcripts localised in viroplasms of rRV-wt infected cells (Fig. 6A), consistent with the roles of viroplasms in supporting the viral replication and assembly. In contrast, no viral RNA transcripts could be detected in the aberrant structures in both wt MA104 or MA-NSP2-mCherry cells infected with the rRV-NSP5/S67A mutant (Fig. 6A). RNA accumulation in viroplasms was instead restored when infecting MA-NSP5 cells that supply wild type NSP5 in *trans* (Fig. 6A-lower panel).

**Figure 6.**
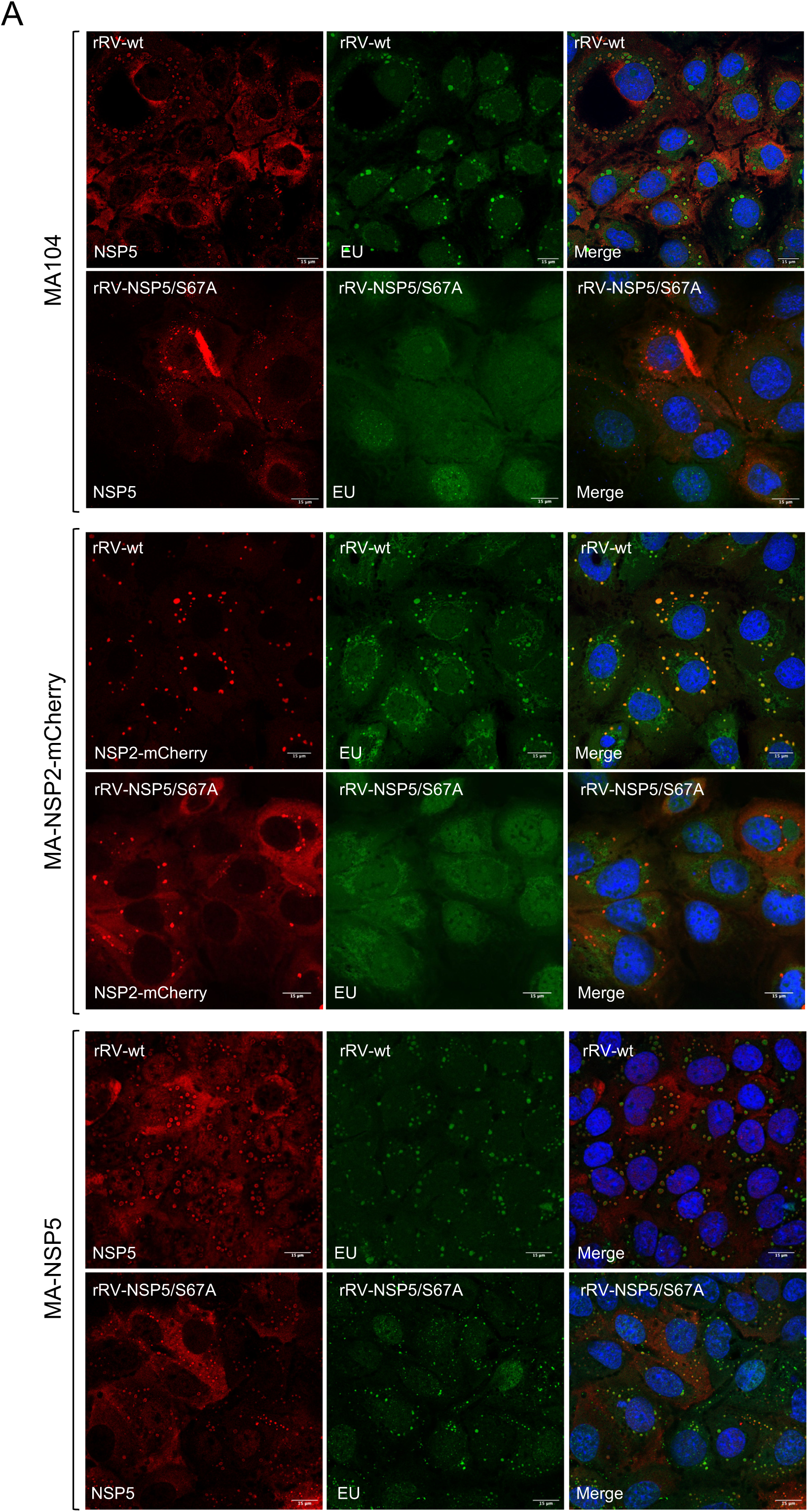
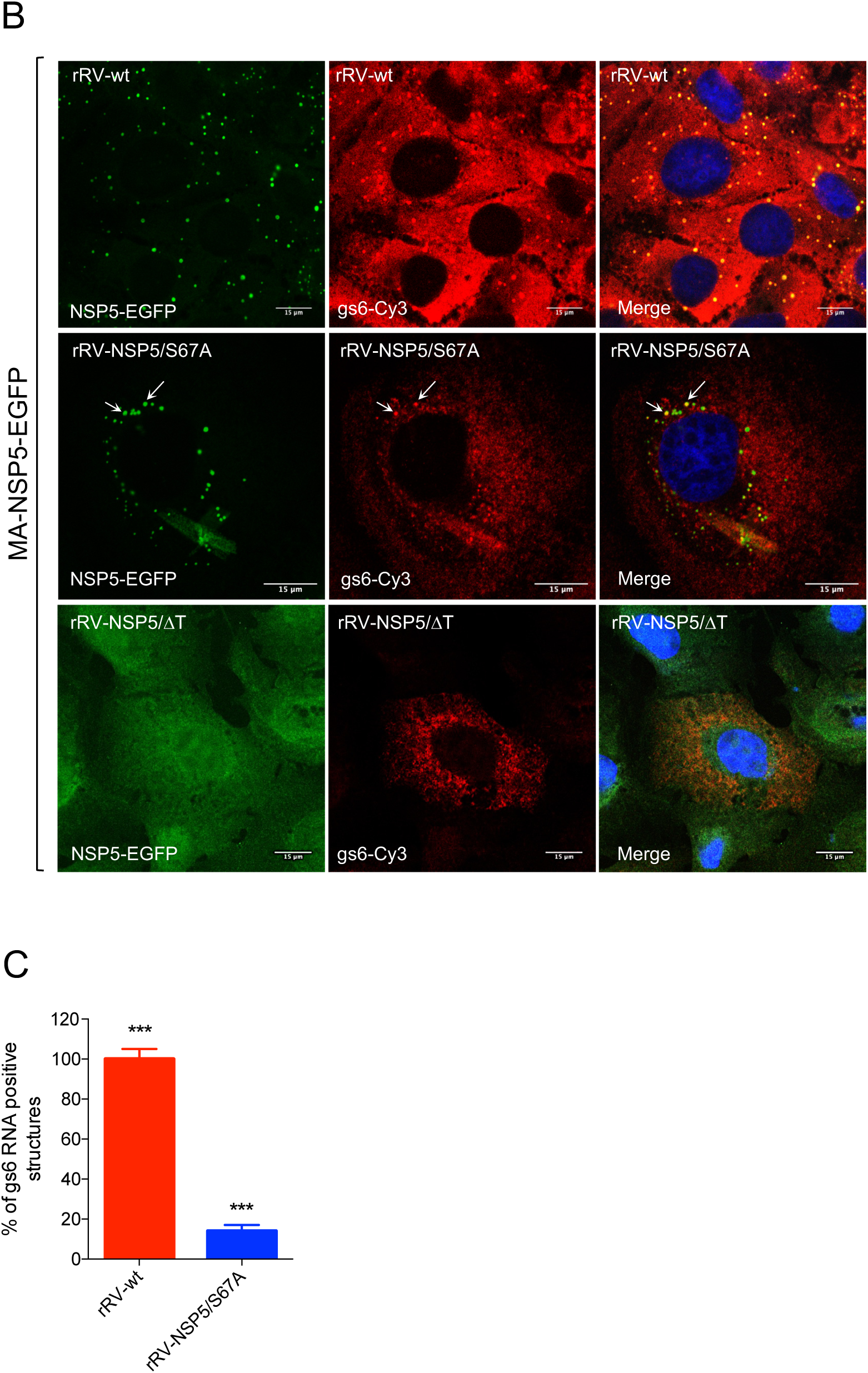
Viral RNA detection in rRV NSP5/S67A infected cells. A) Representative confocal immunofluorescence micrographs of MA104 (upper panel), MA-NSP5 (middle panel) and MA-NSP2-mCherry cells (lower panel) infected with rRV-wt or rRV-NSP5/S67A strains and stained with anti-NSP5 (red, in MA104 and MA-NSP5 cells) and EU (green). B) Single molecule RNA Fluorescence In Situ Hybridisation (smFISH) on MA-NSP5-EGFP cells infected with rRV-wt, rRV-NSP5/S67A or rRV-NSP5/ΔT strains. Viroplasms detected with NSP5-EGFP (green) and viral RNA (probe specific for gs6 was Cy3-conjugated, red). Colocalising viroplasms and RNAs are indicated by white arrows. Scale bar, 15 μm. C) Quantitative analysis of NSP5-EGFP positive structures (viroplasms) for RNA (gs6) in MA-NSP5-EGFP cells infected with rRV-wt or rRV-NSP5/S67A. ***, p < 0.001

Although we did not detect RNA in aberrant structures (or viroplasms) in rRV NSP5/S67A-infected MA104 cells using 5-EU staining, this mutant could still replicate (Fig. 3C). We then explored whether rRV-NSP5/S67A transcripts could accumulate in viroplasms, albeit with much lower efficiency, i.e., beyond the sensitivity limit of 5-EU staining. For this purpose, we used single molecule RNA fluorescence in situ hybridisation (smFISH) to identify the sites of RV transcription. At 10 hpi, abundant gs6 transcripts could be detected in all viroplasms identified in MA-NSP5-EGFP cells infected with the rRV-wt (Fig. 6B). Conversely, the rRV-NSP5/S67A-infected cells had sparse EGFP-tagged structures, but less than 20% of them contain gs6 RNA (Fig. 6B-C). Interestingly, the less frequently occurring rod-like aberrant structures also showed gs6 RNA accumulation, further suggesting that these structures could represent replication-functional organelles.

In contrast, smFISH performed on cells infected with the rRV-NSP5/ΔT showed diffuse distribution of gs6 RNA that did not localise to any structures resembling viroplasms (Fig. 6B), also failing to support the virus genome replication (Fig. 4B).

We then examined the ultrastructures of viroplasms with altered morphologies in the RV mutant rRV-NSP5/S67A using electron microscopy. Upon infection with rRV-wt (Fig. 7A, left panel), multiple membrane-less electron dense inclusions encircled by the well-defined endoplasmatic reticulum (ER) filled with triple-layered particles were present in cells. At late infection points (10 hpi), filled with triple-layered particles (TLPs) ER appeared to adopt a more tubular morphology, suggesting a successive step in the virus egress. In contrast, the rRV-NSP5/S67A-infected cells contained only few immature viroplasms that lacked the ER network filled with TLPs (Fig. 7A, right panel). Only few immature particles containing transient lipid membranes could be identified in cells infected with rRV-NSP5/S67A mutant (Fig. 7A, left panel). Furthermore, the observed immature viroplasms also appeared to be less-electron dense, likely due to their lower RNA composition and, subsequently, decreased number of available phosphate groups that bind UO^2+^ ions during the EM staining procedure. Together, these data strongly support the role of NSP5 phosphorylation in maintaining the viral RNA production and genome replication in viroplasms.

**Figure 7.**
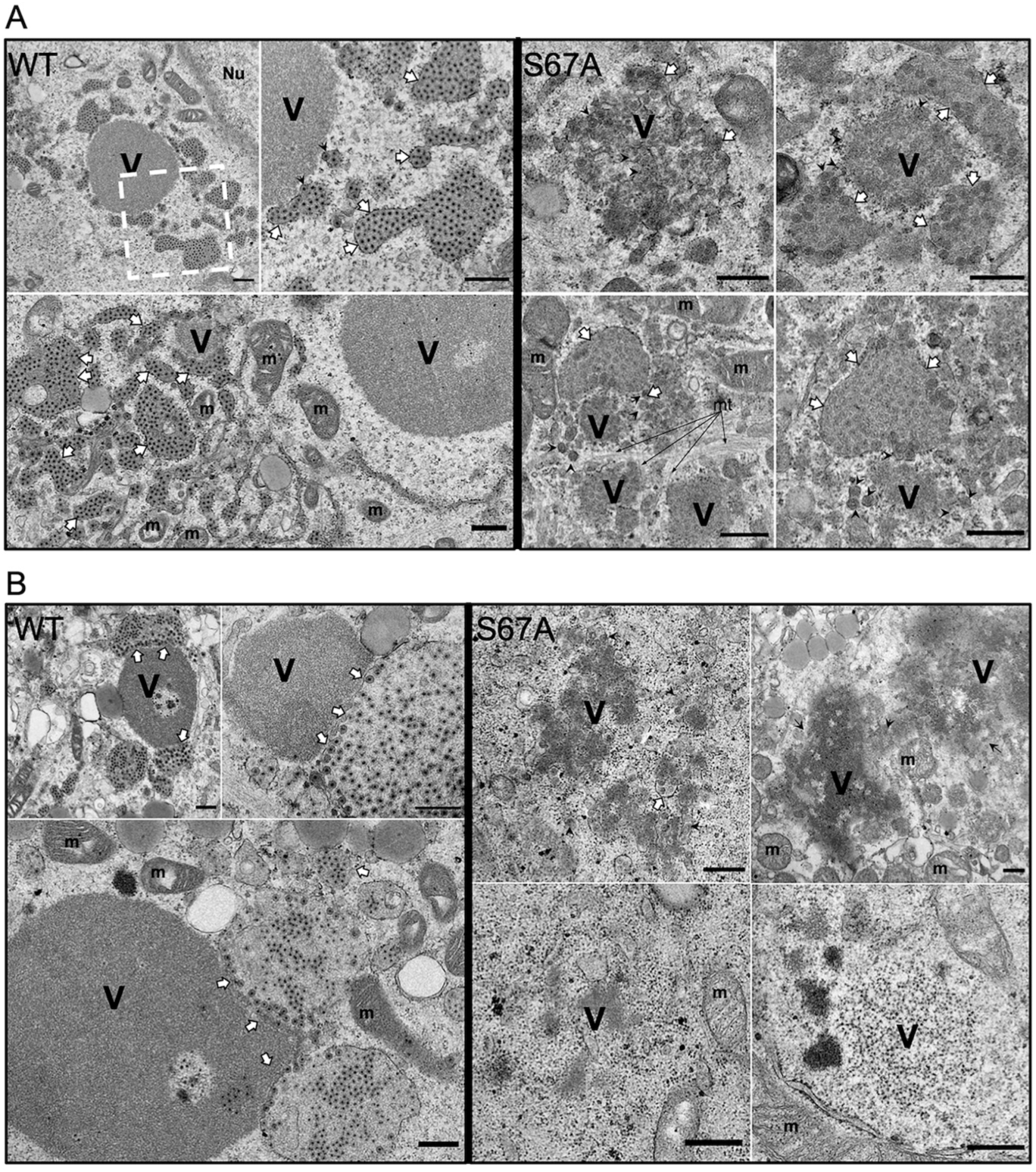
Electron microscopy of cells infected with rRV-NSP5/S67A. High– definition electron micrographs of MA104 (A) and MA-Δ3 (B) cells infected with rRV-wt (left panel) and rRV-NSP5/S67A (right panel) (MOI, 75 FFU/cell). At 10 hpi, cells were fixed with glutaraldehyde and processed for transmission electron microscopy. V, viroplasm; m, mitochondria; mt, microtubule-bundles; Nu, nucleus. The white open box indicates the immediate right magnified image. White arrows indicate the endoplasmatic reticulum surrounding viroplasms; black arrowheads indicate viral particles with an envelope. Scale bar, 500 nm.

#### The mechanism of NSP5 hyper-phosphorylation

We have previously proposed a model of the hierarchical NSP5 hyperphosphorylation associated with the assembly of viroplasms that involves a three-step mechanism: i) initial interaction of non-phosphorylated NSP5 with NSP2 (or VP2); ii) phosphorylation of Ser67 by CK1α. This step does not take place when NSP5 is expressed alone; iii) hyper-phosphorylation of NSP5 triggered by Ser67 phosphorylation that requires the 18 AA long C-terminal tail (24).

Here, we have investigated the phosphorylation mechanism of NSP5 during RV infection using a number of NSP5 phosphorylation-negative rRV strains and MA104-derived stable transfectant cell lines (Table 1). We demonstrated that despite the presence of Ser67, deletion mutant NSP5/ΔT was not phosphorylated and failed to form viroplasms. We have previously shown that co-expression of NSP5/ΔT was also unable to trigger the phosphorylation cascade of NSP5/S67A, while other NSP5 mutants referred hereafter as activators of phosphorylation, e.g., NSP5/Δ3 did (24). Interestingly, upon co-infection with two rRVs NSP5/S67A and NSP5/ΔT, both mutated NSP5 variants were not phosphorylated (Fig. 8A) (8, 19, 24, 26). This result was supported further by infecting the MA104 transfectant cell line stably expressing NSP5-ΔT (MA-ΔT) with the rRV-NSP5/S67A strain (Table 1 and Fig. 8B, lanes 3-4). The NSP5-Δ176-180 mutant was also unable to induce phosphorylation of NSP5/S67A following the co-infection with the two rRVs, despite both containing Ser67 and the C-terminal tail. This result suggests that the ‘activator’ NSP5 needs to be also hyper-phosphorylated (Fig. 8C).

**Figure 8.**
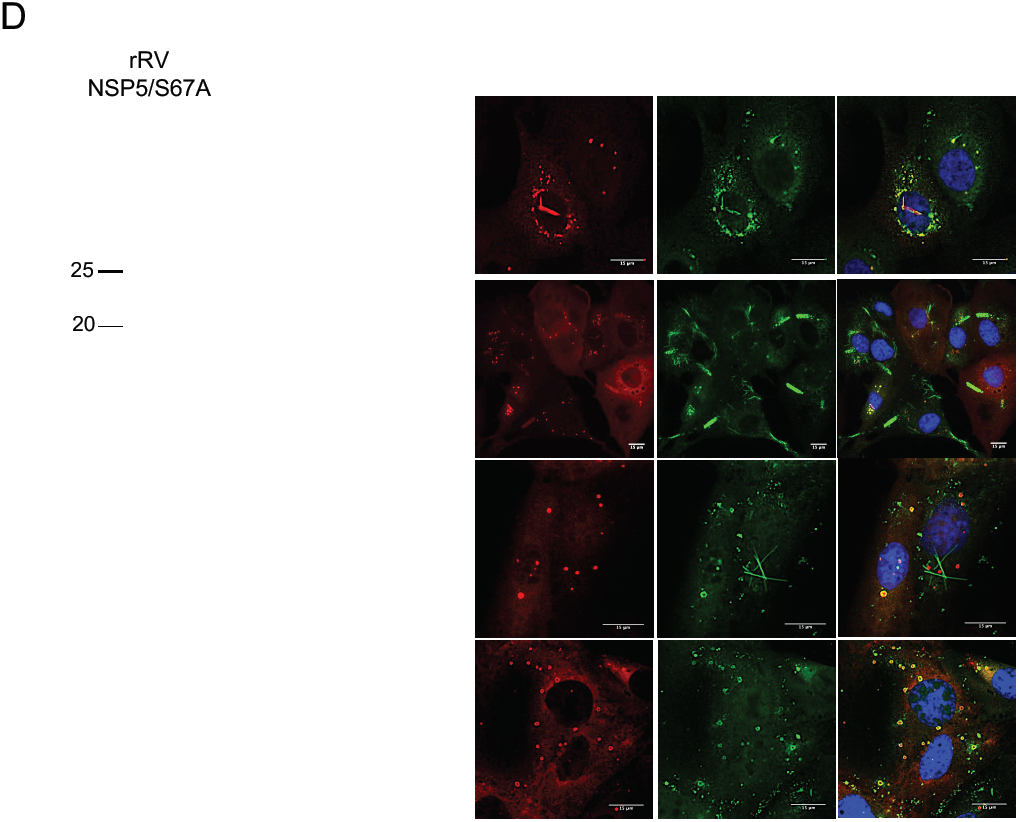
Phosphorylation of NSP5/S67A. A) Western blot of extracts of MA104 cells infected with rRV-NSP5/S67A or co-infected with rRV-NSP5/S67A and rRV-NSP5/ΔT. Blue arrowhead indicates NSP5/S67A. The faster migrating band corresponds to NSP5-ΔT. B) Western blot of MA104, MA-ΔT and MA-Δ3 cells infected with rRV-NSP5/S67A. Blue arrowhead indicates NSP5/S67A. C) Western blot of extracts of MA104 cells co-infected with rRV-NSP5/S67A and rRV-NSP5/Δ176-180. Blue arrowhead indicates NSP5/S67A. D) λ-Ppase treatment of extracts of MA-Δ3 cells infected with the rRV-NSP5/S67A strain. Filled and open arrowheads indicate de-phosphorylated NSP5-S67A and NSP5-Δ3, respectively. E) Representative confocal immunofluorescence micrographs of MA104, MA-ΔT and MA-Δ3 cells infected with rRV-NSP5/S67A. Cells were stained with the indicated antibodies. Scale bar, 15 μm. F) Yield of infectious virus of rRV-NSP5/S67A grown in MA-NSP5, MA104, MA-ΔT and MA-Δ3 cells at 24 hpi.*, *p* < 0.05; **, *p* < 0.01.

In MA-ΔT cells infected with rRV-NSP5/S67A mutant strain aberrant viroplasms were produced (Fig. 8E), similar to those seen in wt MA104 cells, which did not support virus replication compared to the wt virus (Fig. 8F).

In contrast to NSP5/ΔT, the NSP5 deletion mutant lacking amino acids 80-130, (NSP5/Δ3) becomes hyper-phosphorylated when expressed alone and can function as an activator of the NSP5/S67A phosphorylation (19, 24). We therefore asked whether an MA104 stable cell line expressing the deletion mutant NSP5/Δ3 (MA-Δ3) and infected with the rRV-NSP5/S67A strain was able to trigger hyperphosphorylation of NSP5/S67A and as a consequence, sustain replication of the mutant rRV strain. As shown in Fig. 8B (lanes 5-6) NSP5/S67A was hyper-phosphorylated in presence of NSP5/Δ3 mutant and confirmed by the λPpase treatment (Fig. 8D), although it did not completely rescue the phosphorylation pattern of NSP5 normally observed in rRV-wt infection. When loading large amounts of NSP5/S67A, we have occasionally observed a second faint band with reduced mobility, as in Fig. 8B, lane 2 (indicated with *). Interestingly, regular round-shaped structures resembling viroplasms, containing NSP5, NSP2 and VP2 with peripheral localisation of VP6, were recovered in these cells infected with rRV-NSP5/S67A, and yet viral replication was nevertheless impaired (Fig. 8E-F). Consistently, the electron microscopy images showed structures of aberrant viroplasms similar to those obtained in MA104 wt cells (Fig. 7B, right panel).

The NSP5 phosphorylation negative mutants of the two other rRVs (NSP5/ΔT and NSP5/Δ176-180), did not undergo hyper-phosphorylation in MA-ΔT or MA-Δ3 cell lines (Figs. 9A-B). In both cases, viroplasms formation and virus replication were not rescued (Figs. 9C-D and 9E-F). Interestingly, apparently normal round-shaped structures were observed in MA-Δ3 cells, which, however, recruited significantly less VP6 (Fig. 9C-D). In addition, the previously observed VP6 spiky structures were not detected in these cells (Figs. 9C-D). Similar results were obtained with the rRV-NSP5/KO virus strain, which in MA-Δ3 cells did not replicate and also showed spherical structures that contained NSP5 and NSP2, but not VP6 (Fig. 10A-C).

**Figure 9.**
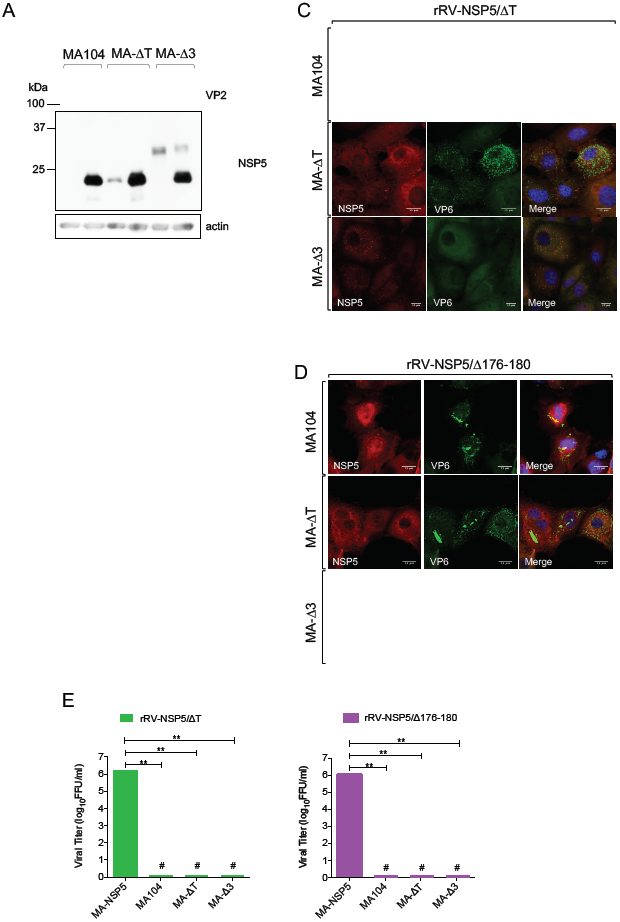
Phosphorylation of NSP5/-ΔT and NSP5/Δ176-180 and viroplasm formation. Western blots (A, B) and confocal immunofluorescence micrographs (C, D) of MA104, MA-ΔT and MA-Δ3 cells infected with rRV-NSP5/ΔT (A, C) or with rRV-NSP5/Δ176-180 (B, D). Blue arrowhead indicates NSP5/ΔT and NSP5/Δ176-180, respectively. Open arrowheads indicate NSP5-Δ3 and NSP5-ΔT. Cells were stained with the indicated antibodies and DAPI. Scale bar, 15 μm. E) Single step growth of rRV-NSP5/ΔT (left) and rRV-NSP5/Δ176-180 (right) in MA-NSP5, MA104, MA-ΔT and MA-Δ3 cells, as indicated. The experiment was terminated at 24 hpi. **, *p* < 0.01; #, Viral titer < 300 FFU/ml.

**Figure 10.**
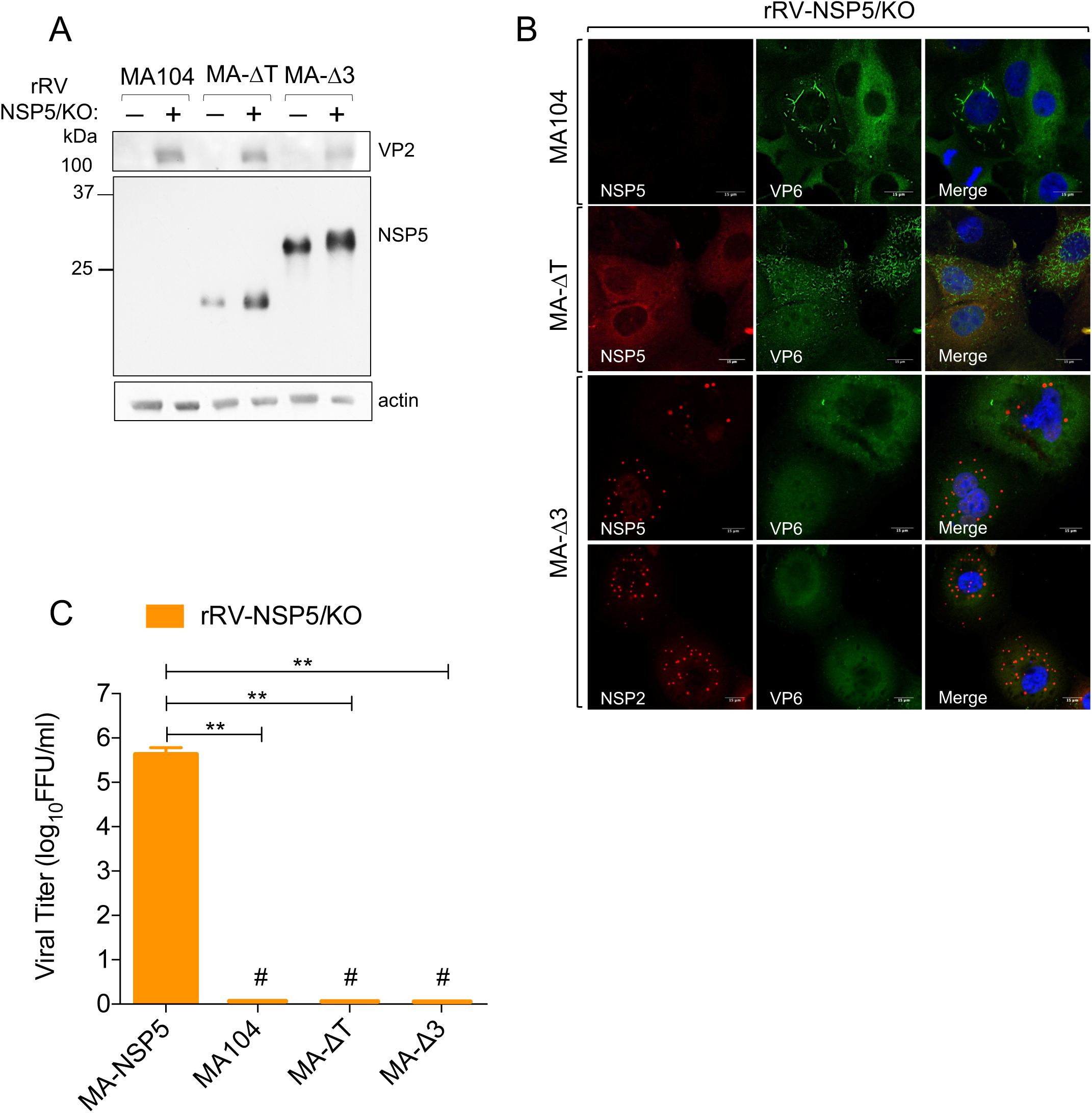
rRV NSP5/KO in infected cells. Western blot (A) and confocal immunofluorescence (B) of MA104, MA-ΔT and MA-Δ3 cells infected with the rRV-NSP5/KO strain. Cells were stained with the indicated antibodies and DAPI. Scale bar, 15 μm. C) Single step growth of rRV-NSP5/KO in MA-NSP5, MA104, MA-ΔT and MA-Δ3 cells, as indicated. The experiment was terminated at 24 hpi. **, *p* < 0.01; #, Viral titer < 300 FFU/ml.

Taken together, our data support a model of NSP5 hyper-phosphorylation, which absolutely requires the presence of the C-terminal tail, and the AA residues 176-180. Furthermore, NSP5 hyperphosphorylation requires Ser67 phosphorylation to initiate the phosphorylation cascade, thus playing a key role in the assembly of replication-competent viral factories.

## DISCUSSION

Rotaviruses replicate within cytoplasmic viral factories, or viroplasms. Most RV assembly intermediates, i.e., single layered particles (cores) and double-layered particles (DLPs) are primarily concentrated in viroplasms. Following budding of DLPs into the lumen of the endoplasmatic reticulum, the immature particles acquire a transient envelope, as well as the outer capsid proteins VP4 and VP7, resulting in a mature triple-layered virion. Moreover, downregulation of expression of the most abundant viroplasm-forming proteins NSP5 and NSP2 severely impacts on the formation of viroplasms and production of virus progeny (13, 15, 25, 34). In light of these observations, viroplasms have long been recognised as essential compartments supporting the RV replication.

Of the two non-structural proteins involved in viroplasm assembly, NSP5 appears to play a crucial role by potentially providing a scaffold that allows for recruitment of additional viral proteins. Only when NSP5 is co-expressed with NSP2 and/or VP2, these proteins assemble into the viroplasm-like structures (VLS), which are also capable to recruit additional structural proteins including VP1, VP2 and VP6 (8, 10). Given these observations, we hypothesised that complete removal of NSP5 would be lethal for RV replication. Using a modified reverse genetics system for rotaviruses (29), here we provide the first direct evidence of the essential role of NSP5 in viroplasm formation and viral replication. In order to characterise replication-deficient NSP5-negative mutants, we have established a trans-complementing system that provides NSP5 to the virus both transiently in BHK-T7 cells, and stably in NSP5-producing MA104 cell line (MA-NSP5), thus enabling facile isolation of rRVs lacking functional NSP5. Using this approach, we have demonstrated that NSP5-deficient rRV was unable to form viroplasms and replicate in the wt MA104 cells, while the viroplasm formation and viral replication were efficiently rescued in the trans-complementing MA-NSP5 cell line. Interestingly, the rRVs generated using this method also failed to incorporate dsRNA originating from the NSP5-encoding mRNA lacking the 5’ and 3’ untranslated regions (UTRs), further suggesting the essential roles of UTRs for genome packaging in RVs (37, 38).

NSP5 hyper-phosphorylation has been previously implicated in the regulation of NSP5 assembly into viroplasms. This phosphorylation, however, requires the interaction of NSP5 with, either NSP2 or VP2, as NSP5 is not phosphorylated when expressed alone (8, 22, 23). Previous studies suggest that activation of NSP5 hyper-phosphorylation may require a conformational change that leads to its efficient hyper-phosphorylation via a positive feedback loop mechanism (8, 23, 24). Two regions comprising the N-terminal amino acids 1-33 (region 1) and the central region amino acids 81-130 (region 3) have been reported to prevent NSP5 phosphorylation in the absence of other viral proteins, while the 18 amino acids long C-terminal tail was found to be essential for its phosphorylation (8, 23, 24).

Here, we have shown that all three rRV mutants S67A, ΔT and Δ176-180 expressing the phosphorylation-negative NSP5 variants were unable to form round-shaped viroplasms upon infection of MA104 cells. Interestingly, further analysis of these mutants reveals some key differences between each NSP5 variant. While rRV-NSP5/S67A strain formed aberrant structures resembling viroplasms that poorly support RV replication, this variant was still capable of producing the infectious progeny, in contrast to the two other rRV mutant strains. Interestingly, the phenotype observed with the NSP5 mutant S67A was essentially the same as the one previously reported with the wt virus infecting MA104 cells silenced for expression of CK1α, which is involved in phosphorylating Ser67 and initiating the hyper-phosphorylation cascade (24, 25). It has recently been shown that CK1α is also involved in phosphorylating NSP2, controlling the formation of rotavirus viral factories (39). Our data obtained with the rRV-NSP5/S67A mutant strongly suggest that the lack of NSP5 hyper-phosphorylation determines both the morphogenesis of viroplasms and their capacity to support RV genome replication. We cannot rule out the role of NSP2 phosphorylation in assembly of viroplasms since NSP2 is also likely to be phosphorylated by CK1α upon infection of the rRV-NSP5/S67A strain. Despite the formation of aberrant structures resembling viroplasms, the amount of RNA produced within those structures in rRV-NSP5/S67A-infected MA104 cells was practically below the detection limit of the 5-EU labelling, while the RNA replication was fully rescued in the trans-complementing MA-NSP5 cell line (Fig. 5). It is unlikely that the viral mRNAs produced in MA104 wt cells infected with the rRV-NSP5/S67A strain was degraded faster, as most of the ssRNA synthesised is a consequence of the secondary round of transcription from the newly made dsRNA-containing particles. Indeed, very low amounts of dsRNA were detected during the infection of MA104 cells. smFISH results confirmed the presence of small amounts of RV (+)ssRNA in some of these aberrant structures. This result is consistent with the finding that the rRV-NSP5/S67A strain did replicate, albeit at much lower levels than the rRV-wt. Thus, these structures are likely to sustain virus replication with decreased efficiency, which was further confirmed by the electron microscopy analysis of MA104 cells infected with the rRV-NSP5/S67A strain.

The important role of NSP5 hyper-phosphorylation was further supported by the results obtained with the two phosphorylation-negative mutant strains NSP5/ΔT and NSP5/Δ176-180 that possess Ser67 and yet failed to form viroplasms in MA104 cells. Surprisingly, the Δ176-180 mutant, containing the C-terminal tail has also failed to form viroplasms in MA104 cells, despite the absence of Ser or Thr within the chosen 176-180 region. Both phosphorylation-negative mutants tested did not replicate in MA104 cells and we could not detect any structures containing viral RNA.

Using the NSP5 mutant strains described above and the established stable transfectant MA104 cells we were able to investigate the molecular mechanism that leads to NSP5 hyperphosphorylation. We showed that the NSP5/S67A mutant from the rRV was indeed hyperphosphorylated, albeit not completely, when infecting MA-Δ3 cells, restoring the round-shape morphology of the structures resembling viroplasms with a complete absence of the aberrant structures observed in MA104 cells. This finding strongly suggests that impairment of NSP5 phosphorylation is the direct cause of the formation of the aberrant structures in the cytosol of the infected cell. Despite the fact that these structures appeared morphologically similar to the classical round-shaped viroplasms in MA-Δ3 cells, the presence of VP6 around these structures was only observed with the hyperphosphorylated NSP5/S67A mutant, in contrast to the other NSP5 phosphorylation-negative mutants. One possibility is that, during RV infection, accumulation of VP6 in round-shaped viroplasms requires a full-length hyperphosphorylated NSP5, as well as phosphorylation of multiple serine residues likely by CK2 (17, 26). Moreover, the round-shaped structures found in MA-Δ3 cells infected with rRV-NSP5/KO failed to contain VP6.

The observed failure to rescue the replication of the rRV-NSP5/S67A strain to the wt levels in MA-Δ3 cells could be the consequence of the incomplete recovery of the complex pattern of phosphorylated isoforms of wt NSP5. This suggests that some intermediate isoforms might be important for the formation of fully functional replication-competent viroplasms.

We propose a model of the complex hierarchical mechanism of NSP5 hyper-phosphorylation during RV infection (Fig. 11). It involves (a) interaction of NSP5 with either NSP2 or VP2, required to (b) make Ser67 available for CK1α phosphorylation. This initial step is then sequentially completed (c) by CK2-mediated phosphorylation of other serines to generate the NSP5 ‘activator’, during a step dependent (d) on the interaction of the Ser67-phosphorylated molecules with the non-phosphorylated partners in the NSP5 oligomeric complexes. Alternatively (c’), oligomers could be formed before the activation step. NSP5 interactions mediated by the carboxy-terminal tails T result in (e) substrate activation and a fully hyperphosphorylated NSP5. This process leads to (f) the assembly of viroplasms scaffolds containing NSP2, and (g) recruitment of VP6, as well as the other viroplasmic proteins, VP1, VP2, VP3 to assemble replication-competent viroplasms.

**Figure 11.**
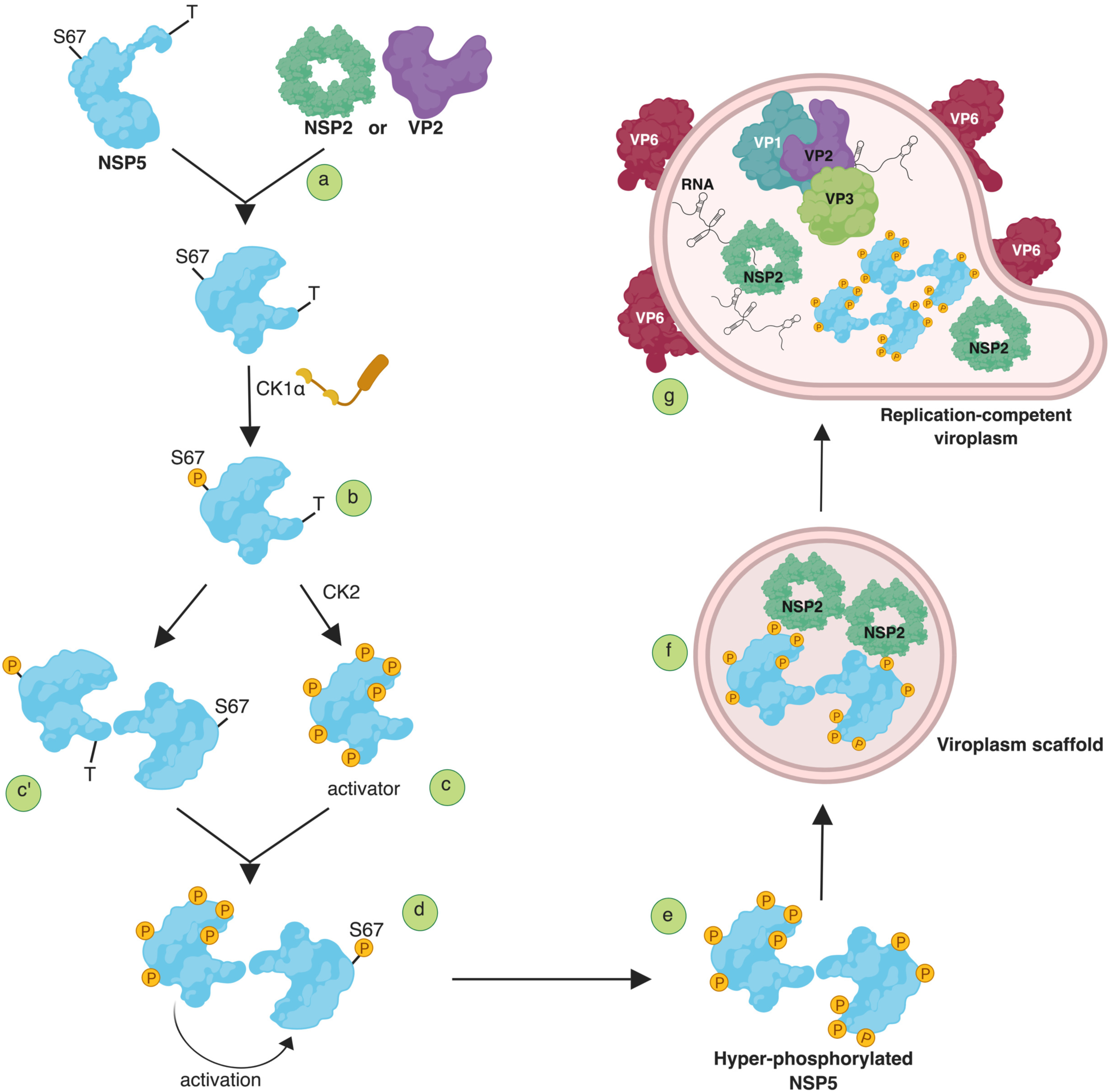
Model of NSP5 hyperphosphorylation and assembly of replication-competent viroplasms. (***a***) Interaction of non-phosphorylated NSP5 with either NSP2 or VP2 is required to (***b***) induce conformational changes that make S67 available for CK1α phosphorylation (P). This initial step of the cascade is then (***c***) sequentially completed by phosphorylation by CK2 of other residues including serines in domain 4, to generate the NSP5 activator or, (***c’***) a step of interaction with non-phosphorylated molecule precedes the involvement of CK2. (***d***) NSP5 interactions to form dimers/multimers are mediated by the carboxy-terminal tails (T). (***e***) The primed (S67-phosphorylated) activator molecules result in substrate activation and fully hyper-phosphorylated NSP5. (***f***) Assembly of viroplasms scaffolds containing NSP2 serve to (***g***) recruit the other viroplasmic components, VP1, VP2, VP3 and VP6 to assemble replication-competent viroplasms. P; phosphorylated amino acid. *This image was created using BioRender.com.

The hierarchical phosphorylation of proteins appears to be a common mechanism regulating many cellular processes. In mammalian cells, hierarchical phosphorylation has been described for β-catenin, in which Ser 45 phosphorylated by CK1α primes it for hyperphosphorylation by glycogen synthase kinase-3 (GSK-3) (40, 41), which triggers its ubiquitination and proteasomal degradation (42). A similar phosphorylation mechanism has recently been described for a non-structural protein NS5A of the Hepatitis C Virus (HCV), with the hyper-phosphorylation cascade primed by the initial phosphorylation of Serine 225 by CK1α, and the subsequent phosphorylation of neighbouring residues involving other kinases (43, 44). Moreover, the NS5A phosphorylation was shown to play a key role in controlling the establishment of replication complexes during HCV infection (45). Similarly, a number of other non-structural viral proteins have been shown to undergo multiple phosphorylation events during virus infection, suggesting that this complex post-translational modifications play a pivotal role in orchestrating the assembly of replication-competent viral factories (46–48).

## ACKNOWLEDGEMENTS

We thank John T. Patton, Asha Ann Philipp and Jin Dai (Indiana University, Bloomington, USA), Yuta Kanai and Takeshi Kobayashi (Osaka University, Osaka) and Albie Van Dijik (North-West University, South Africa) for helpful discussions on the establishment of the reverse genetics system. We are grateful to Naoto Ito for providing BHK-T7 cells.

